# Within-finger maps of tactile and nociceptive input in the human parietal cortex

**DOI:** 10.1101/599167

**Authors:** Flavia Mancini, Martin I. Sereno, Min-Ho Lee, Giandomenico D Iannetti, Irene Tracey

## Abstract

The spatial representation of nociceptive input in the human parietal cortex is not fully understood. For instance, it is known that the primary somatosensory cortex (S1) contains a representation of nociceptive-selective input to different fingers, but it is unclear whether S1 subdivisions contain finer-grained, within-finger maps of nociceptive input. It is also unknown whether within-finger maps of somatosensory input exist in intraparietal regions. Therefore, we used high-field 7T functional MRI to reveal within-finger maps of nociceptive and tactile inputs in the human parietal cortex. Although we replicated the previous findings of between-finger maps of nociceptive input spanning S1 subdivisions, we found weak and inconsistent evidence for within-finger maps of nociceptive input in S1 subdivisions. In the same subjects, we found mirrored, within-finger maps of tactile input in areas 3a, 3b, and 1. Importantly, we discovered a within-finger map of nociceptive input, but not of tactile input, in the human intraparietal area 1 (hIP1). In conclusion, our study indicates that the spatial representation of nociceptive input in the parietal cortex partly differs from that of tactile input and reports the discovery of a within-finger map of nociceptive input in hIP1.

**New & Noteworthy:** We report the discovery of a fine-grained map of nociceptive input to finger segments in the human intraparietal area 1.

## Introduction

Human parietal areas contain topographic representations of innocuous somatosensory input (Sereno and Huang 2014), which support spatial processing of tactile input, sensorimotor transformations and goal-directed behaviour. Although these regions represent predominantly non-nociceptive somatosensory information (Frot et al. 2013), they are also the target of spinothalamic projections conveying nociceptive information (Dum et al. 2009; Gingold et al. 1991; Kenshalo et al. 2000; Kenshalo et al. 1988; Kenshalo and Isensee 1983; Schmahmann and Pandya 1990). However, the precise spatial organization of nociceptive responses in the parietal cortex is not clear.

In particular, each of the four cytoarchitectonic areas (3a, 3b, 1, and 2) within the primary somatosensory cortex (S1) contains a complete representation of mechanical, non-nociceptive input to the hand and fingers, both along the *medial-to-lateral axis* (e.g., *between* finger representation) and along the *anterior-to-posterior axis* (*within* finger representation) (Kaas et al. 1979; Merzenich et al. 1978). The within-finger tactile map in each area of S1 mirrors the representation in the adjacent area; the base-to-tip finger representation is posterior-to-anterior in area 3b but anterior-to-posterior in the adjacent area 1 (Kaas et al. 1979; Merzenich et al. 1978; Sanchez-Panchuelo et al. 2012).

Previous studies have shown that the human S1 contains somatotopic maps of nociceptive inputs along its medio-lateral axis, spanning different cytoarchitectonic subdivisions: not only the broad somatotopies for hand, face, and foot territories (Andersson et al. 1997; Bingel et al. 2004), but also the between-fingers maps of nociceptive input (Mancini et al. 2012). It is unknown whether and how nociceptive input is mapped along the anterior-to-posterior axis of S1. Spinothalamic projections also target areas in the intra-parietal sulcus (IPS) and superior parietal lobule (SPL) (Schmahmann and Pandya 1990). It is unknown whether IPS contains a representation of somatosensory input to different finger segments. Therefore, we used high-field functional MRI (fMRI) to determine whether within-finger maps of nociceptive input exist in the parietal cortex, if they exist, to evaluate whether these maps are co-aligned to within-finger maps of tactile input.

## Methods

### Participants

Fourteen healthy and right-handed subjects (3 women) aged 21–32 years (mean ± SD: 24.64 ± 3.39) participated in the study, after having given written informed consent. All experimental procedures were approved by the Health Research Authority (REC reference: 11/SC/0249; IRAS projectID: 80049). Twelve participants undertook a within-finger mapping protocol. Additionally, two participants undertook a between-finger mapping protocol for replication of previous results obtained at lower magnetic field strengths (Mancini et al. 2012; Mancini et al. 2018) and for validating our scanning and analyses methods.

### Experimental design

#### Within-finger mapping

Twelve participants attended two scanning sessions, each lasting 75 minutes. The sessions were identical except for the modality of the somatosensory stimulus: mechanical and non-nociceptive stimulation of Aβ fibers in one session, nociceptive-selective stimulation of Aδ/C fibers in the other session, in counterbalanced order across participants (see Supplemental Data for a description of the stimuli used: https://github.com/flamancini/parietalmaps/blob/master/supplemental_data_JNP19.pdf). Participants laid inside the scanner bore with their right hand open and supine, kept firmly in place with the aid of foam pads and a cushion. The stimulation was a travelling wave of somatosensory stimuli sweeping through the proximal segments of digits 2, 3, 4 and 5, the middle segments of digits 2-5, and the distal segments of digits 2-5. The different digits were stimulated in random order and only the proximal-to-distal stimulation was given in sequential order. To correct for systematic regional variations in the shape of the hemodynamic response function, we interleaved progressions in two directions (proximal-to-distal; distal-to-proximal). The first progression was counterbalanced among participants. For both modalities, 12 cycles of 42.67s trains of stimuli (interstimulus interval, 2.13s) were administered in each of the three runs. Each run both started and ended with 20s of rest. However, painful sensations elicited by high-energy laser pulses can typically last for more than a second. One participant withdrew from the study before completing the full protocol and was not included in any analysis.

#### Between-finger mapping

To validate our scanning and analyses methods, we mapped the between-finger representation of tactile stimuli in one participant and that of nociceptive stimuli in another participant. All procedures were identical to the ‘within-finger mapping’ except for the spatial progression of the stimuli. The stimulation was a travelling wave sweeping through random locations in d2, d3, d4, and d5. Each digit was stimulated sequentially, whereas the different segments of each digit were stimulated randomly (but covering the same surface of the within-finger mapping protocol). Again, we interleaved progressions in two directions (d2-to-d5; d5-to-d2).

### MRI data acquisition

MRI data were acquired on a Siemens Magnetom 7 Tesla MRI scanner with a 32-channel head coil. Blood oxygenation level dependent (BOLD) fMRI was acquired using multislice gradient echoplanar imaging, with axial slices centred on the typical location of hand maps in S1 (voxel size 1.7mm isotropic, 281 volumes/run, 30 axial slices, flip angle 65º, TR 2000ms, TE 26ms, GRAPPA acceleration factor PE 2, bandwidth 1370Hz/Px, echo spacing 0.81ms). The slice coverage of our echoplanar images was centred on the hand-knob in S1 and included most of the parietal lobe and premotor cortex, but excluded most of the cingulate cortex and insular cortex. Task-fMRI data were collected during 6 runs (3 runs per session; 1 session per modality). The first 4 measurements of each functional run were discarded from all subsequent analyses.

We also collected a structural MP2RAGE scan (voxel size 1mm isotropic, 192 slices per slab, flip angle 1 4°, TR 5000ms, TI 1 900ms, TE 2.84ms, GRAPPA acceleration factor PE 3, bandwidth 240Hz/Px, echo spacing 7 ms) and a gradient field map (voxel size 2mm isotropic, 64 slices, flip angle 39°, TR 620ms, TE 1 4.08ms, TE 2 5.1ms, bandwidth 734Hz/Px). We used the FreeSurfer program ‘recon-all’ (https://surfer.nmr.mgh.harvard.edu/) to reconstruct the cortical surface from the T1 structural image, after correcting for image intensity non-uniformity using the AFNI program ‘3dUniformize’ (https://afni.nimh.nih.gov) and non-brain removal with 3dSkullStrip.

### FMRI analyses

We performed two parallel sets of analyses: Generalized Linear Model analyses (GLM) in order to study the magnitude of the cortical response; phase mapping analyses in order to characterise the spatial organisation of parietal responses.

#### GLM analyses

The purpose of these analyses was to determine the magnitude of brain responses to tactile stimuli and nociceptive stimuli in S1 and IPS, irrespective of the somatotopic organization of these responses. Data processing was carried out using standard GLM analyses in FSL (www.fmrib.ox.ac.uk/fsl; supplemental data https://github.com/flamancini/parietalmaps/blob/master/supplemental_data_JNP19.pdf). fMRI data were subjected to exclusion in cases of visible spin history motion artefact as a result of sharp motion during more than one scan sessions (1 mm of absolute mean displacement in fewer than five volumes), as in previous studies (Kolasinski et al. 2016a; Kolasinski et al. 2016b); this happened in two participants, which were then excluded from further analyses. Thus, nine participants were included in further analyses. The two participants who undertook the between-finger stimulation paradigm were not included in univariate analyses, because we did not collect data from both modalities in each of the two participants. Task-based statistical parametric maps were computed separately for the tactile condition and nociceptive condition versus resting baseline. At group level, we used FLAME1+2 to test: (1) whether activations in the tactile condition were greater than baseline; (2) whether activations in the nociceptive condition were greater than baseline; (3) whether activations in the tactile condition were different than in the nociceptive condition. Z statistic images were thresholded non-parametrically using clusters determined by Z>2.3 and a corrected cluster significance threshold of p=0.05.

#### Phase-encoded analyses

The purpose of these analyses was to determine whether any parietal region showed a preferential response to the spatial frequency of the periodic finger stimulation; voxels sensitive to a given spatial frequency of stimulation respond maximally when the stimulus passes through the preferred location and decay as the stimulus moves away (Chen et al. 2017). Surface-based analyses were performed on unsmoothed data as previously described (Mancini et al. 2018) (see Supplemental Data for a detailed description of the analyses https://github.com/flamancini/parietalmaps/blob/master/supplemental_data_JNP19.pdf).

## Results

### Magnitude of cortical responses

At the group level, both tactile and nociceptive stimuli to the fingers activated clusters in S1, superior parietal lobule, and anterior intra-parietal cortex in the hemisphere contralateral to hand stimulation (Fig. 1, Supplemental Table I and II https://github.com/flamancini/parietalmaps/blob/master/supplemental_data_JNP19.pdf). Tactile stimuli significantly activated a S1 region in the hemisphere ipsilateral to hand stimulation (Fig. 1A). No brain region was activated more by nociceptive stimuli than tactile stimuli. In contrast, tactile stimuli elicited greater responses in S1 than nociceptive stimuli, both in terms of response amplitude and spatial extent (Fig 1C). This difference is unlikely to be mediated by trivial factors, such as differences of perceived intensity or saliency of the two stimulus modalities. First, although S1 responses were lower for nociceptive stimuli than tactile stimuli, the average perceived intensity of pinprick pain was higher than the average perceived intensity of touch (t_23_=4.06, p=0.0004; mean ‘touch’ intensity rating: 4.18, SD across participants: 1.03; mean ‘pinprick pain’ intensity rating: 6.24, SD across participants: 1.48). Second, the energy of the laser stimuli was set as the highest the participants agreed to endure for 3 runs of ~8 minutes of stimulation. Third, the laser energy was adjusted before each run to ensure that pain intensity remained at the same level of the first run (mean rating variability across runs: 0.38).

**Figure 1.**
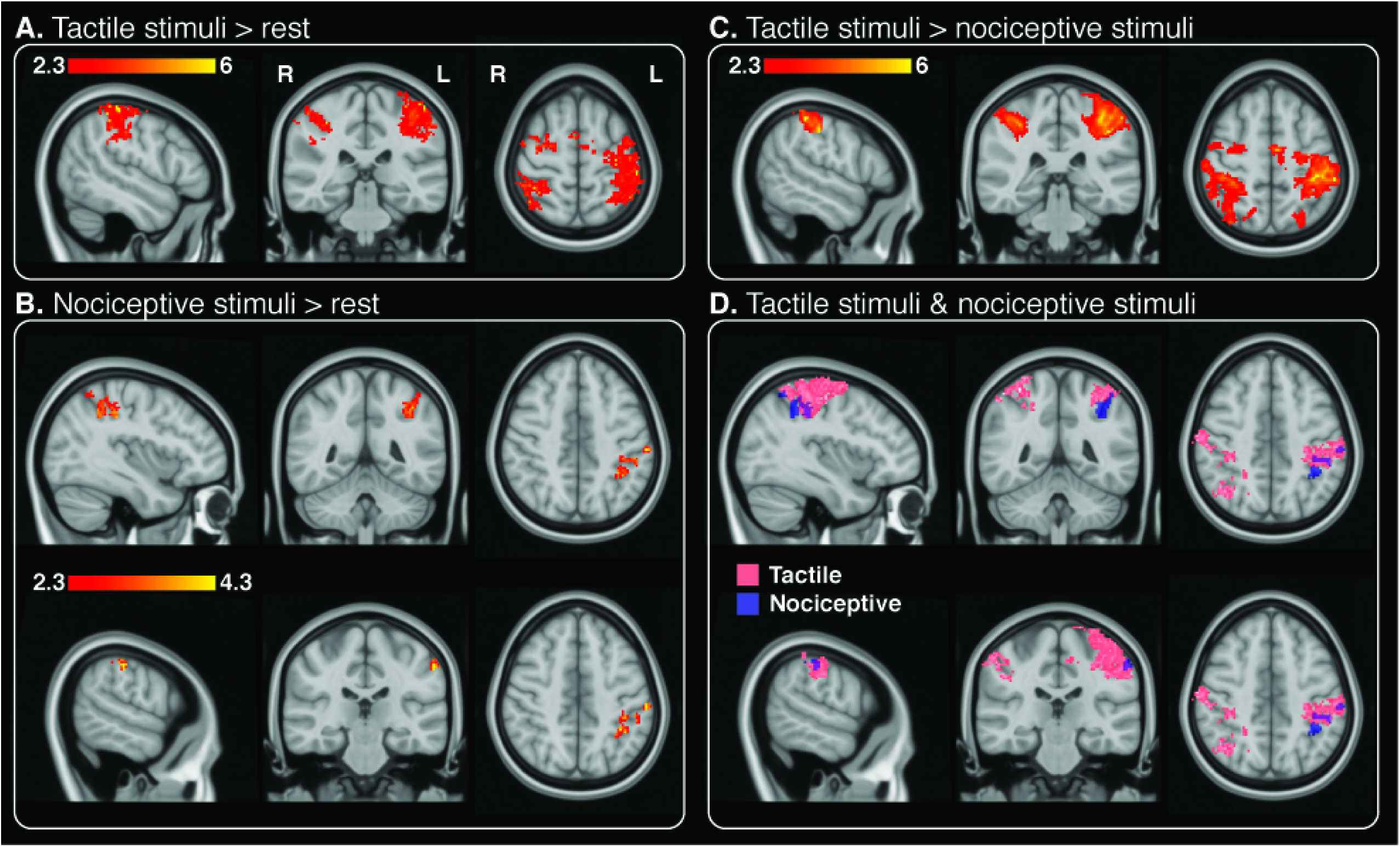
Cortical responses to tactile (**A**) and nociceptive stimuli (**B**), as quantified using GLM analyses. Panel **C** shows regions more activated by tactile stimuli than nociceptive stimuli, whereas panel **D** shows the overlay of cortical activity elicited by tactile and nociceptive stimuli. The displayed Z statistics were thresholded using clusters determined by Z>2.3 and a corrected cluster significance threshold of p=0.05.

### Somatotopic organization of cortical responses

First, we validated our 7T mapping protocols by replicating our previous findings of contralateral, between-finger maps of tactile and nociceptive input in S1 (Fig.2). As we already described (Mancini et al. 2012; Mancini et al. 2018), the between-finger map of both tactile and nociceptive stimuli progressed along the lateral-to-medial axis from index finger to little finger. Then, we went on searching for within-finger maps in S1 and IPS. At the group level (Fig.3A-C), we found mirrored maps of tactile input to finger segments across S1 subdivisions, in the hemisphere contralateral to hand stimulation. For reference, the estimated anatomical boundaries of S1 subdivisions were estimated using an automated and probabilistic surface-based parcellation. The ROIs were statistically thresholded and displayed on the inflated cortical surface of an average brain. The tactile finger map progressed, along the anterior-to-posterior axis, from base to fingertip in area 3a. The phase of the map reversed approximately at the border between area 3a and 3b, at the fingertip. We did not observe consistent phase reversals between areas 3b and 1, but we found a significant finger segments map in area 1, that progressed from tip to base of the fingers along the anterior-to-posterior axis (Fig.3B-C).

**Figure 2.**
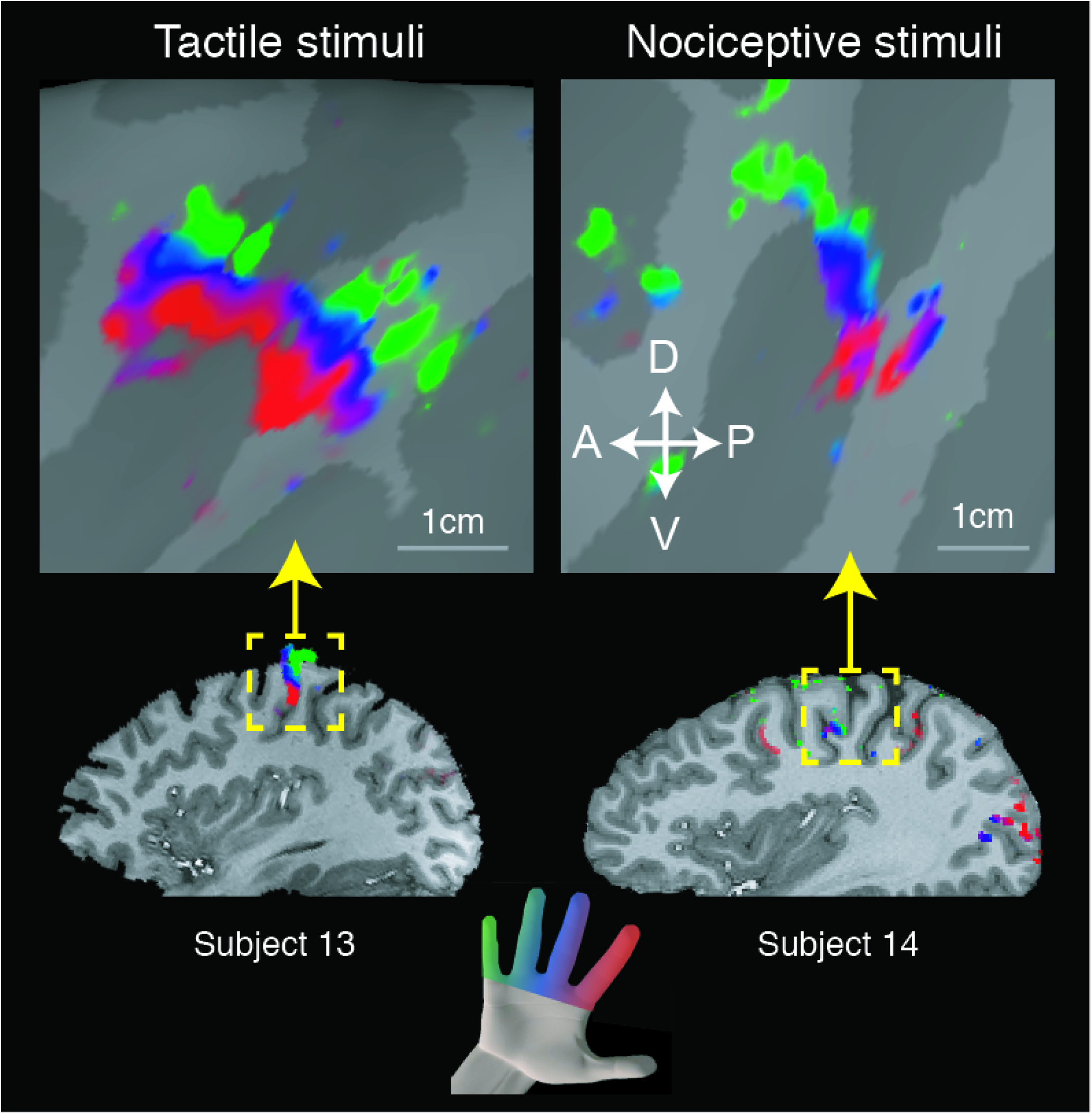
Replication of previous evidence of between-finger maps of tactile stimuli (subject #13) and nociceptive stimuli (subject #14), as revealed by phase-encoded mapping analyses. The F-statistics of the signal at different phases of periodic finger stimulation (a travelling wave sweeping through the different fingers, from the index finger to the little finger) are rendered on the T1 volume and on the inflated cortical surface.

**Figure 3.**
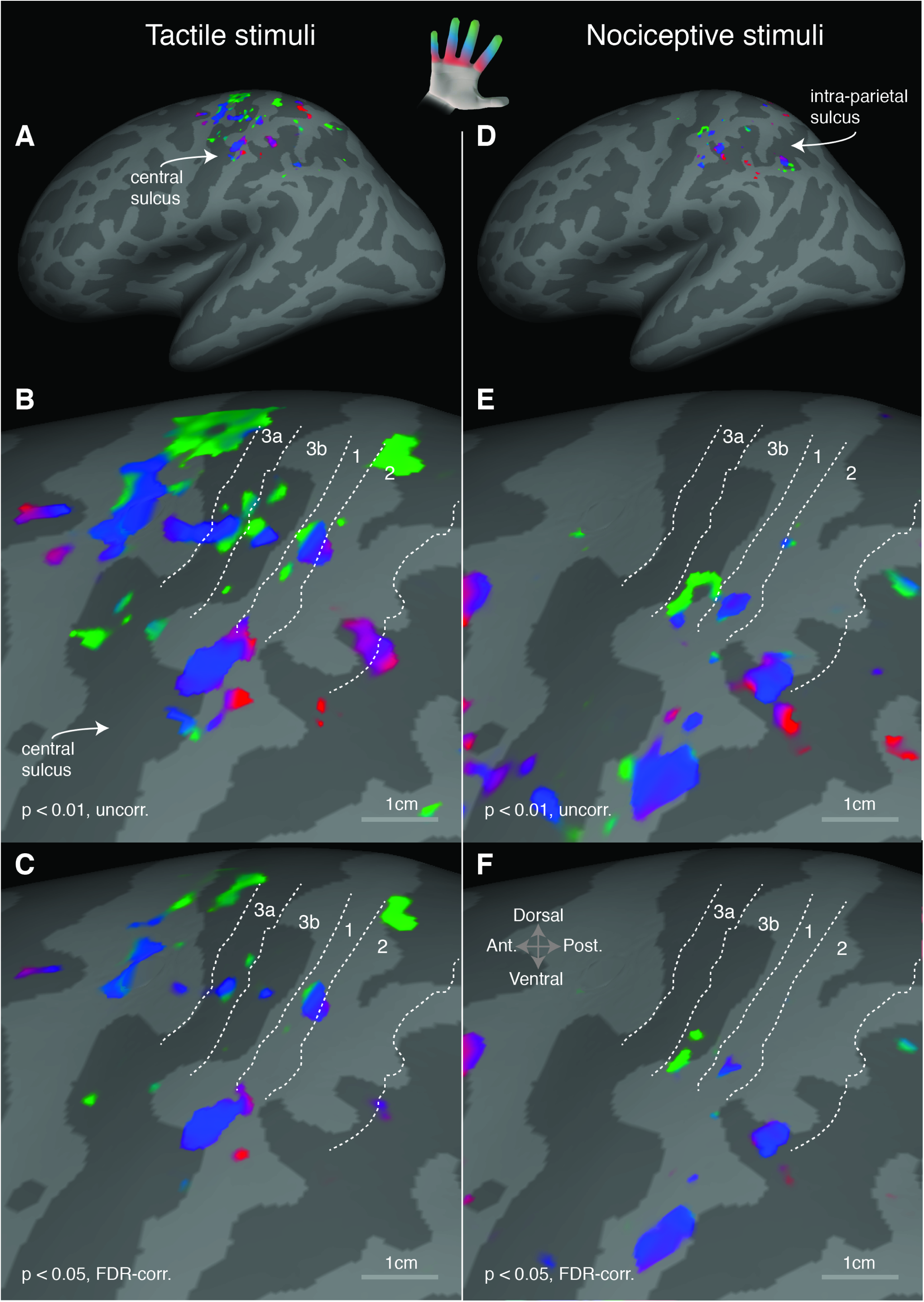
Surface-based, group average of cortical responses elicited by periodic within-finger tactile (**A-C**) and nociceptive (**D-F**) stimulation. The color-coding scheme used is shown on the top of the figure: red = proximal finger segments, blue = medial finger segments, green = distal finger segments. Panels **A, B, D** and **E** display the uncorrected F-statistics, thresholded at p<0.01. Panels **C** and **F** show the F-statistics with an FDR-correction for p-values at level 0.05. Tactile stimuli, but not nociceptive stimuli, exhibit consistent within-finger maps in S1 subdivisions. The F-statistics is rendered onto an inflated average cortical surface (fsaverage) and overlaid the anatomical boundaries between areas 3a, 3b, 1 and 2 (estimated using Freesurfer’s probabilistic surface-based parcellation and thresholded).

Nociceptive stimuli elicited a periodic response in areas 3b and 1 (Fig.3D-E), in a more ventro-lateral region than the one activated by tactile stimuli. The responses to nociceptive stimuli appeared to follow a similar spatial gradient than those to tactile stimuli, at group level. However, these activity clusters barely survived FDR correction (Fig.4F) due to large intra-individual variability. Therefore, we cannot exclude that these within-finger activations in S1 might be spurious. Nociceptive stimuli, but not tactile stimuli, elicited periodic responses in a region in the contralateral human intraparietal area 1 (hIP1) (Choi et al. 2006; Scheperjans et al. 2008), as shown in the group average maps (Fig.4A-C). The centroid of this nociceptive map was located at Talairach coordinates −43, −52, 37 (MNI coordinates −43, −55, 39). In the majority of subjects (Fig.4D), we found significant periodic responses to nociceptive stimulation of finger segments.

**Figure 4.**
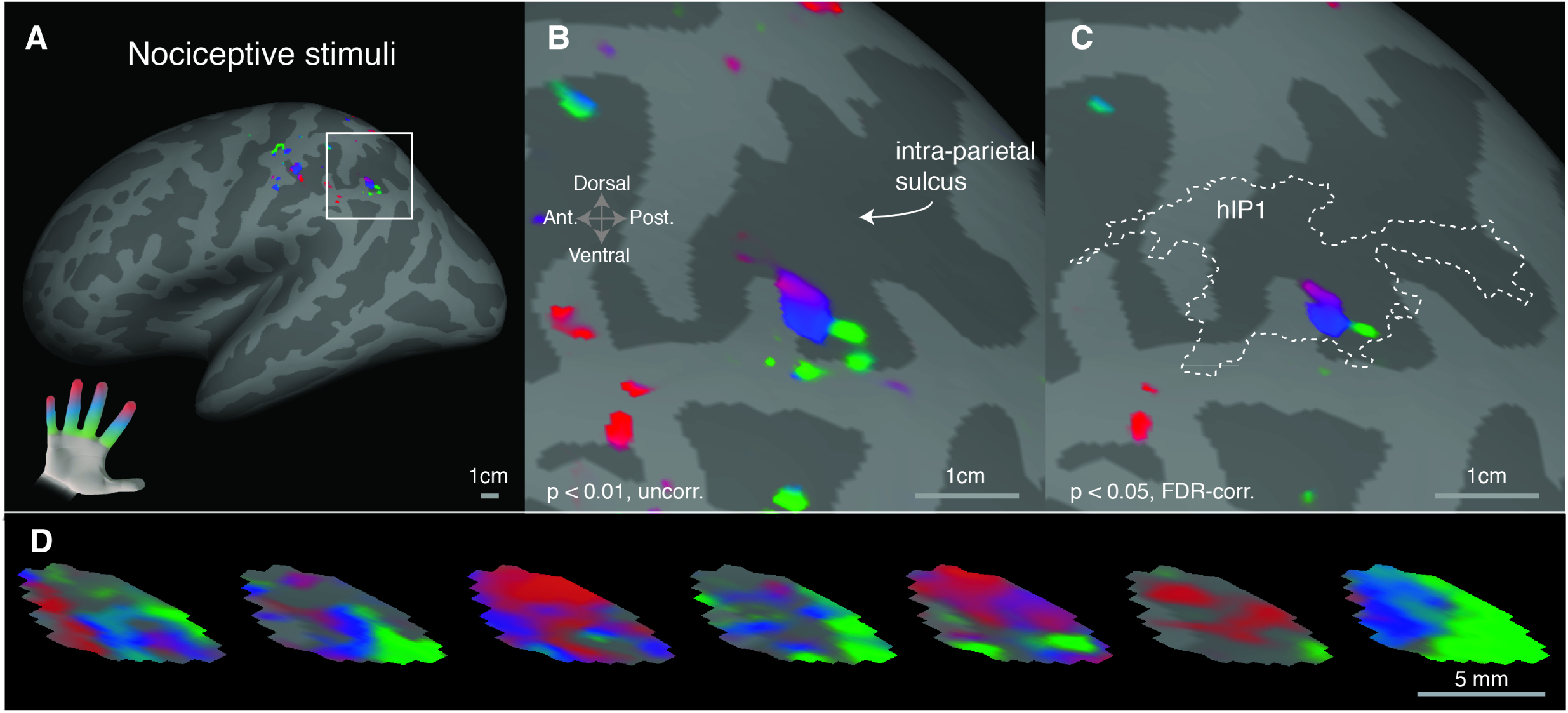
**A-C**. Surface-based, group average of within-finger maps of nociceptive stimuli (n=9) in human intraparietal area 1 (hIP1). The color-coding scheme is as follows: red = proximal finger segments, blue = medial finger segments, green = distal finger segments. Panels **A-B** display the uncorrected F-statistics, thresholded at p<0.01, whereas panel C shows the F-statistics with an FDR-correction for p-values at level 0.05. For reference, in panel C the anatomical boundaries of area hIP1 are displayed (as derived from the probabilistic Juelich Histological Atlas and thresholded at 25%). **D**. Single-subject, within-finger maps of nociceptive input (one map per subject; the two subjects that did not show any significant response in this region are not displayed). The surface-based mask used to display the single-subject data corresponds to the boundaries of the group-average, within-finger map of nociceptive input displayed in panel B.

## Discussion

Our study indicates that the somatotopic representation of nociceptive input in the parietal cortex differs from that of tactile input. In particular, we did not find robust evidence for the existence of within-finger maps of nociceptive-selective stimuli across S1 cytoarchitectonic subdivisions, but we were able to successfully detect within-finger maps of tactile input in Brodmann areas 3a, 3b, and 1, in the hemisphere contralateral to hand stimulation (Fig.3). However, we confirmed the existence of between-finger maps of nociceptive input in the medial-to-lateral axis of S1 (Fig.2), in line with previous reports (Mancini et al. 2012; Mancini et al. 2018). Although nociceptive stimuli did not elicit consistent BOLD responses in contralateral (left) S1, they consistently activated areas in a region in the left intra-parietal area 1 (hIP1), which did not display a significant periodic response to tactile stimulation (Fig.3-4).

### Within-finger maps of tactile input in S1

Only a few fMRI studies have investigated the within-finger somatotopy of mechanical input in the human S1 (Blankenburg et al. 2003; Overduin and Servos 2004; Sanchez-Panchuelo et al. 2012; Schweisfurth et al. 2011). To date, only one of these studies (Sanchez-Panchuelo et al. 2012) has been able to reliably identify the border of map reversals in individual subjects. In monkeys, the locations of reversals of within-finger somatotopic maps identified from electrophysiology and anatomy have consistently been found to coincide with cytoarchitectonic borders (Kaas et al. 1979; Merzenich et al. 1978; Nelson et al. 1980).

The within-fingers somatotopy of tactile input reported here is largely consistent with that reported by previous work; we found that the most anterior reversal was around the fingertips, at the estimated border between areas 3a and 3b (Fig. 4B-C). Although we did not observe, at the group level, a map reversal between areas 3b and 1, we found that the spatial arrangement of the tactile within-finger map in area 1 followed the one expected on the basis of previous work (Sanchez-Panchuelo et al. 2012) (Fig. 4B-C). At the group level, we did not observe a within-finger map in area 2.

### Nociceptive responses in S1

It is clear from animal and human studies that a subset of S1 neurons responds to nociceptive input, but their functional relevance and spatial organization remains controversial (Vierck et al. 2013). Studies in primates support a possible nociceptive role for areas 3a and 1 (Kenshalo et al. 2000; Tommerdahl et al. 1996; Whitsel et al. 2009). In humans, nociceptive responses originating in areas 1–2 have been suggested from MEG and subdural EEG data (Inui et al. 2003; Kanda et al. 2000; Ogino et al. 2005; Ploner et al. 1999), but the involvement of area 3b in nociception remains controversial (Baumgartner et al. 2011; Kanda et al. 2000; Valeriani et al. 2004). Intracerebral recordings in the human S1 have shown that responses specific to nociceptive heat in area 3b were sparser and much less frequent than responses to non-nociceptive input (Frot et al. 2013). Our study suggests that the density of nociceptive units might be sparse or heterogeneously distributed in S1 along the anterior-to-posterior axis, and may account for weak and inconsistent within-finger representations.

### Within-finger maps of nociceptive input in hIP1

Our study reveals a novel within-finger map of nociceptive input in hIP1 (Choi et al. 2006). The map was significant at both group and individual-subject level in the majority of participants (Fig.4). This sub-region of hIP1, or any other intraparietal subregion, was not significantly activated by tactile stimuli (Fig.3A). To our knowledge, no previous study has demonstrated the existence of within-finger maps of somatosensory input in the intraparietal cortex, possibly because the parietal somatotopy of the hand is both spatially and temporally variable across individuals (Chen et al. 2017).

Nociceptive stimuli have frequently been reported to activate intra-parietal posterior parietal regions (Duerden and Albanese 2011; Tracey and Mantyh 2007) and these activations are usually interpreted as reflecting spatial attention (Lobanov et al. 2013; Oshiro et al. 2009; Oshiro et al. 2007; Peyron et al. 1999). It is possible that we were able to detect a within-finger map in hIP1 only in response to nociceptive input, but not tactile input, because nociceptive stimuli are inherently more salient (being aversive) than non-painful tactile stimuli (Legrain et al. 2011). Indeed, previous work has shown that the anterior intraparietal cortex is sensitive to bottom-up attention driven by stimulus saliency (Geng and Mangun 2009). Future studies are required to confirm the functional significance of the within-finger map of nociceptive input in hIP1.

## Acknowledgments

This project has received funding from the Wellcome Trust (Strategic Award <u>102645/Z/13/Z</u>). The Wellcome Centre for Integrative Neuroimaging is supported by core funding from the Wellcome Trust (<u>203139/Z/16/Z</u>).

